# Efficient site-specific editing of the *C. elegans* genome

**DOI:** 10.1101/007344

**Authors:** Donglei Zhang, Michael Glotzer

**Author notes:** Correspondence should be addressed to M.G.

## Abstract

In just two years, genome editing with the CRISPR-associated endonuclease Cas9 has transformed genetic analysis in conventional and emerging model organisms. The efficiency of this method varies among systems and continues to be optimized. Numerous strategies have been reported for editing the *C. elegans* genome. To date, these strategies do not provide a simple, rapid and inexpensive means to introduce and isolate arbitrary point mutants. Here, we report a strategy with all three of these desirable properties. It utilizes oligonucleotides as donor templates for homology-dependent repair and visible markers that are edited in parallel that markedly reduce the number of animals that must be molecularly screened in order to isolate mutants that lack visible phenotypes.

## INTRODUCTION

Fast and affordable genome editing by the CRISPR-associated endonuclease Cas9 has been demonstrated in diverse organisms ^1^. The initial applications of this method involved introduction of small insertions and deletions (indels) to create loss of function mutations and the addition of epitope tags, such as GFP ^2,3^. CRISPR/Cas9 has also been deployed in *C. elegans* by a number of investigators (see ^4^ for review). Existing strategies for isolation of edited genomes involve screening for edited genomes by easily scored phenotypes, genotyping of individual worms, or embedding of a selectable marker in the donor template. The incorporation of a selectable marker simplifies the isolation of edited genomes, but it requires construction of large, variably complex, repair templates, requires a second step to trigger loss of the selectable marker, and ultimately results in a residual LoxP site ^5^. Engineering single nucleotide substitutions for structure-function studies can require even more complex repair templates. Alternative strategies that eliminate the need for large repair templates have been recently described, but the edited genomes are difficult to isolate when the sequence change does not result in an easily scored phenotype ^6^.

The ability to rapidly incorporate single nucleotide changes at native genomic loci would greatly improve the power and reliability of structure-function analysis of proteins of interest. Therefore we sought to develop a rapid, reliable, and inexpensive method to incorporate and identify small changes at arbitrary loci. To that end we used programmable CRISPR/Cas9, single-stranded DNA oligonucleotide (ODN) repair templates, and a second, independent and easily scored repair event to facilitate identification of CRISPR-edited genomes. With this strategy, mutations of interest can be readily isolated in < 2 weeks.

## RESULTS AND DISCUSSION

### Short ODNs are efficient donors for Cas9 trigged homologous recombination

We first tested the efficiency of CRISPR using ODNs as repair templates. We made two changes to the donor templates as compared to the recent report demonstrating that ODNs can function as repair templates in *C. elegans*^6^. Specifically, we use shorter ODNs (60 nt vs 100 nt, see Table 1 for oligonucleotides used in this study), and, we introduced silent mutations, either in the protospacer adjacent motif (PAM) or in the codons just upstream of the PAM ^7^, in order to render edited genomes resistant to the sgRNA. We used microinjection to provide Cas9, sgRNA and 60-nt ODN donor template into the gonad of young adult nematodes. This approach successfully corrected the *unc-119(ed3)* allele, introduced a dominant *rol-6*^D^ mutation, and corrected a single copy integrated non-fluorescent (NF)-GFP (Figure 1).

**Table 1.**
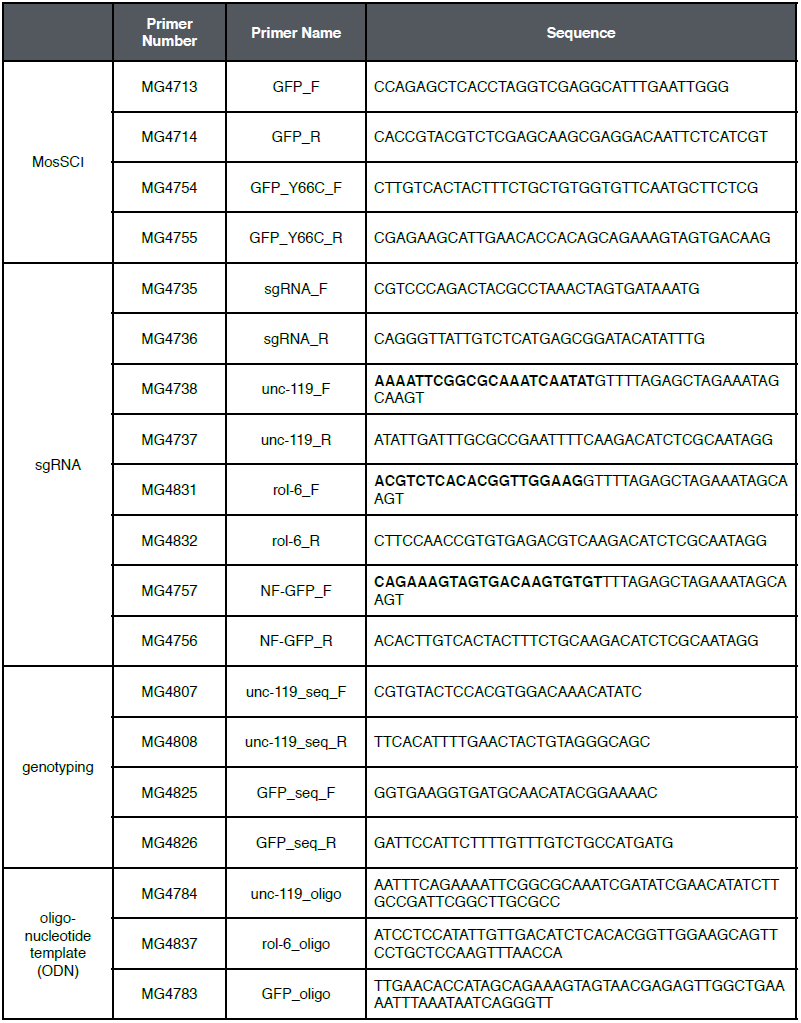
List of oligonucleotides used in this study. Target sequences of sgRNA are highlighted.

**Figure 1.**
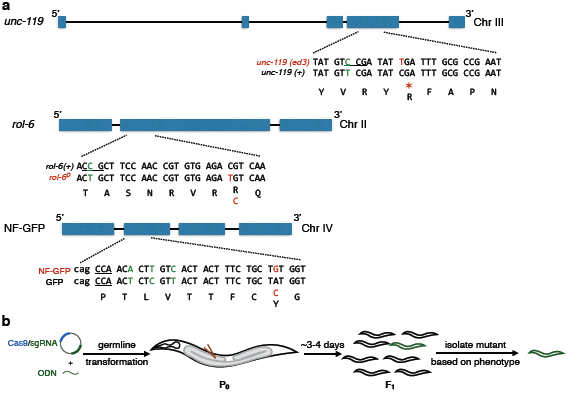
Schematic of target gene editing by CRISPR/Cas9 and single-stranded DNA oligonucleotides (ODNs). (a) Gene editing at *unc-119*, *rol-6*, and NF-GFP loci. In all cases, the desired mutations (red) and additional mutations to abolish the protospacer adjacent motif (PAM) or sgRNA target sequence (green). PAMs are highlighted by underline. Lower case, intron. Uppercase, exon. (b) Schematic for target gene editing by CRISPR/Cas9 system and ODNs (Details in Materials and Methods).

All of these events, being functionally dominant, were readily identified in the F1 generation of the injected worms by visual inspection. Edited genomes were obtained with high efficiency; 15% ∼ 40% of injected worms yielded strains heterozygous for the desired mutations (Figure 2). The frequency of successful editing was higher in healthy strains with large brood sizes (i.e N2, NF-GFP) as compared to rescuing less healthy mutants to wild-type. The frequency of editing events among the total pool of F1 animals was obviously lower, ranging from 0.17% to 0.41% of all F1 animals (Figure 3). This does not serve as a significant barrier to isolating mutants with readily scored dominant phenotypes, but it necessitates extensive molecular screening for mutants that are not easily recognized. With one exception, CRISPR-edited F1s were heterozygous for the desired mutations, and all edited genomes segregated to F2 animals (i.e. the F1s are not somatic mosaics). This suggests that the majority of the double stand break (DSB) and homology directed repair (HDR) events occur in the germline of injected P0s. In the one exceptional case, the animal was heterozygous for an edited genome and a small deletion.

**Figure 2.**
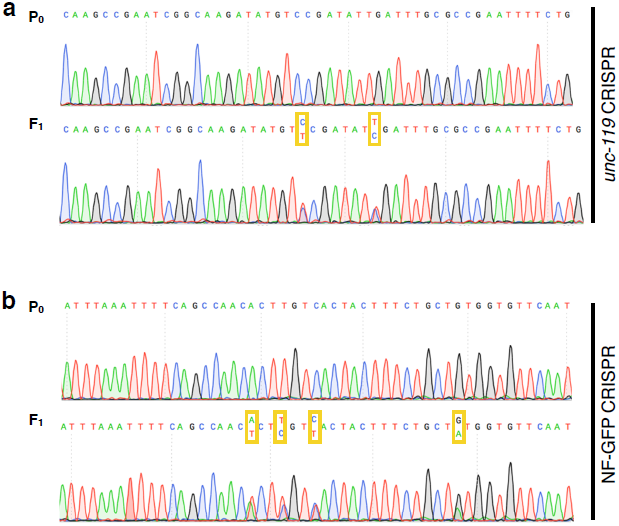
Sequence confirmation of induced mutation in (a) *unc-119(ed3)* and (b) NF-GFP. Nearly all F1 progeny are heterozygous for desired mutations (see Figure 1); 1 of the 48 F1 mutants had both chromosomes edited, one with the desired mutation and the other with a 6bp deletion (see text). Sequence reads near sgRNA target region are shown, and edited nucleotides are boxed.

**Figure 3.**
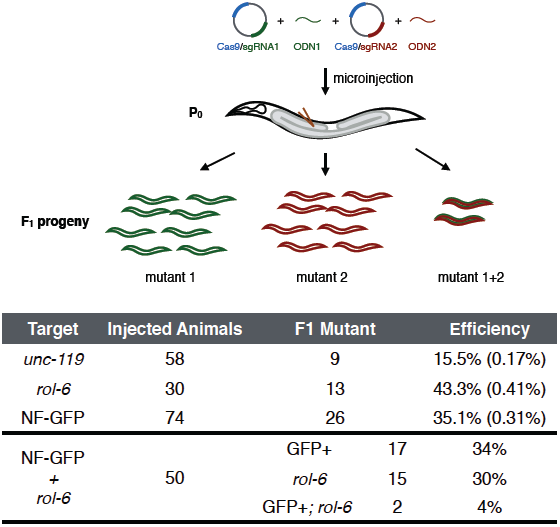
Editing multiple target genes by CRISPR/Cas9 and single-stranded DNA oligonucleotides (ODNs) directed homologous recombination (HR). (a) Schematic for editing two target genes in one-step. (b) Summary of results from individually editing 3 different target genes and coupled editing at NF-GFP and rol-6 loci. Efficiency is calculated as the ratio of F1 mutants to number of injected animals. The percentage of F1 animals with edited genomes is indicated in parentheses.

ODNs as short as 60 nt can serve as repair templates for Cas9/CRISPR guided gene editing. It is thought that HDR requires a minimum length of homology between donor and recipient. For example, in *Drosophila*, HDR requires donors > 0.2-kb donors ^8^. The increased efficiency of 60 nt ODNs may result from the high molar concentrations of short ODNs as compared to plasmid template or PCR-generated dsDNA donor. It is notable that in the case of the *rol-6*^D^ ODN, only 18 nucleotides lie between the introduced mutation and the 3′ end, however, the efficiency of this ODN was similar to that of *unc-119* and NF-GFP. Thus, short ODNs with limited homology to the target are efficient donors for HDR.

### Co-injection markers do not enrich for ODN-based homologous recombination

To facilitate identification of silent mutations, we first tested whether the pool of F1 animals transgenic for a standard co-microinjection marker used for germline transformation were enriched for CRISPR-edited genomes. We injected Cas9, NF-GFP or *unc-119* sgRNAs, corresponding ODNs, and a *Pmyo-3*::mCherry co-injection marker. Although many *Pmyo-3*::mCherry positive animals were produced, none were GFP-positive nor *unc-119*+ (Figure 3) ^6^.

### ODNs serve as donors for multiple targets by a co-CRISPR strategy

Given the demonstrated efficiency of multiplex CRISPR in other systems ^9^, we reasoned that a second CRISPR edited allele might be a more effective transformation marker. We therefore investigated whether alleles with dominant, visible phenotypes, including *unc-119*, *rol-6* and NF-GFP, could facilitate identification of otherwise silent mutations. To examine this approach, we microinjected Cas9, NF-GFP and *rol-6* sgRNAs, and GFP and *rol-6*^D^ ODNs into NF-GFP worms and isolated 17 GFP+, 15 *rol-6*^D^, and 2 GFP+; *rol-6*^D^ strains (Figure 3). While editing events are rare (∼0.25%) in the total pool of F1 animals, 10% of animals edited at one locus are also edited at the second locus, reflecting an enrichment of 40-fold. Thus, by sequencing ∼ 20 strains containing a visible edited marker, one has a high likelihood of isolating a phenotypically silent mutation (a typical workflow is shown in Figure 4). Indeed, this method enabled us to generate a phenotypically silent mutation with a similar efficiency at another site (manuscript in preparation).

**Figure 4.**
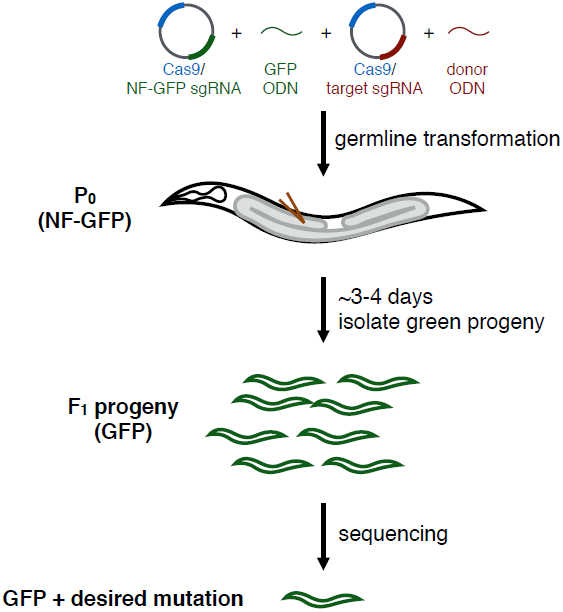
Workflow for generation and identification of phenotypically silent mutants by coCRISPR and ODNs. Cas9/sgRNA and ODN for mutation of interest are microinjected into young adult worms with Cas9/sgRNA and ODN of an easily scored dominant marker (e.g. NF-GFP/GFP). After 3∼4 days, isolate F1 progeny based on the dominant marker (GFP worms, for example). After brooding to maintain line, the gene of interest is amplified from positive F1 progeny to identify those with desired mutation. The whole process can be completed in <2 weeks.

While this manuscript was in preparation, a conceptually similar approach was reported. In particular, the activity of a validated sgRNA that induces a visible phenotype could serve as a marker for a second sgRNA. However, that study used plasmid donor templates, not ODNs. Additionally, HDR events were enriched by selecting for animals transgenic for an extrachromosomal co-injection marker (e.g. *rol-6*^*dm*^) ^10^. In our study, ODN-directed repair events were not enriched in animals expressing analogous extrachromosomal arrays.

## CONCLUSION

Here, we described an improved method to perform genome-editing to introduce single nucleotide changes into *C. elegans*. The use of short, inexpensive ODNs as substrates and easily scored dominant markers for successful genome editing allows inexpensive and rapid identification of phenotypically silent mutants.

## MATERIALS AND METHODS

### Strains

Animals were grown at 20°C on standard nematode growth media (NGM) plates seeded with OP50 *Escherichia coli*. Some strains were provided by the *Caenorhabditis* Genetics Center. The following strains were used: N2, EG8081 *unc-119(ed3) III; oxTi177 IV*, MG827 *mgSi36[Pmyo-2::GFP(Y66C)::let-858 UTR, cb-unc-119(+)] IV; unc-119(ed3) III*, and *unc-119(ed3)*.

### Generation of single copy integrated non-fluorescent (NF)-GFP strain

The single copy integrated non-fluorescent (NF)-GFP strain is generated by the universal mosSCI method developed recently ^11^. To inactivate the chromophore in GFP, we introduced a Y66C substitution. PCR was used to amplify overlapping segments of *Pmyo-2::gfp::let-858 3′UTR* from pPD118.33 (primers: MG4713/MG4755; MG4714/MG4754). The PCR products were inserted into AvrII/XhoI-cleaved pCFJ150 by the SLiCE cloning method ^12^. pCFJ150-*Pmyo-2::NF-GFP::let-858 UTR*, pCFJ104, pMA122, and pCFJ601 were transformed into EG8081 *unc-119(ed3) III; oxTi177 IV*. *Pmyo-2::NF-GFP::let-858 UTR* transgenic worms were identified as described ^11^, and confirmed by sequencing.

### Cas9 target site selection

Cas9 target sites were analyzed with the Zhang lab CRISPR design tool (http://crispr.mit.edu/). Target sequences closest to sites of desired nucleotide changes were selected.

### Cas9/sgRNA plasmid construction

Cas9/sgRNA plasmids were derived from pDD162 vector ^5^. sgRNA target sequences were generated by overlapping PCR using pDD162 as PCR template and the appropriate primers from Table 1. Overlapping PCR products were inserted into pDD162 linearized with SpeI/BsrBI by SLiCE ^12^. All sgRNA constructs were verified by sequencing.

### Microinjection

Microinjection was performed by injecting DNA mixture into gonad arms of young gravid hermaphrodites (P_0_). Generally, the injection mixture consists of Cas9/sgRNA plasmids and oligonucleotide templates (ODNs). Injected P_0_s were maintained at 25°C individually until F_1_s with desired phenotypes were isolated.

The final concentrations of plasmids and ODNs:

> *unc-119* editing experiment: Cas9/*unc-119* sgRNA vector at 50 ng/µl, and *unc-119* oligonucleotide at 50 ng/µl.
>
> rol-6 editing experiment: Cas9/*rol-6* sgRNA vector at 60 ng/µl, and *rol-6* oligonucleotide at 50 ng/µl.
>
> NF-GFP editing experiment: Cas9/NG-GFP sgRNA vector at 50 ng/µl, and GFP oligonucleotide at 50 ng/µl.
>
> NF-GFP/rol-6 coCRISPR experiment: Cas9/NG-GFP sgRNA vector at 60 ng/µl, Cas9/*rol-6* sgRNA vector at 60 ng/µl, GFP oligonucleotide at 50 ng/µl, and *rol-6* oligonucleotide at 50 ng/µl.

In experiments using coinjection marker, we used pCFJ104 (*Pmyo-3*::mCherry) at 5ng/µl.

### Genotyping and identification of edited worms with desired mutations

F1 progeny with desired phenotypes were isolated onto individual plates 3∼4 days after injection of the P_0_. F1 animals were allowed to lay eggs, then the adult was lysed for analysis. The genomic region covering introduced mutations was amplified by PCR and sequenced.

## AUTHOR CONTRIBUTIONS

D.Z. and M.G. co-designed the project and wrote the manuscript. D.Z. performed all experiments.

## ACKNOWLEDGMENTS

D.Z. was supported by the Chicago Fellows Program of the University of Chicago and NIH grant R01GM085087 to M.G.

